# The ataxin-1 interactome reveals direct connection with multiple disrupted nuclear transport pathways

**DOI:** 10.1101/438523

**Authors:** Sunyuan Zhang, Nicholas A. Williamson, David A. Jans, Marie A. Bogoyevitch

**Affiliations:** Cell Signalling Research Laboratories, Department of Biochemistry and Molecular Biology, and University of Melbourne, Parkville, Victoria 3010, Australia; Bio21 Molecular Science and Biotechnology Institute, University of Melbourne, Parkville, Victoria 3010, Australia; Nuclear Signalling Lab., Department of Biochemistry and Molecular Biology, Monash University, Clayton, Victoria 3800, Australia

## Abstract

The expanded polyglutamine (polyQ) tract form of ataxin-1 drives disease progression in spinocerebellar ataxia type 1 (SCA1). Although polyQ-ataxin-1 is known to form distinctive intranuclear bodies, the cellular pathways and functions it influences remain poorly understood. Here, we identify direct and proximal partners constituting the interactome of ataxin-1[85Q] in Neuro-2a cells. Pathways analyses indicate a significant enrichment of essential nuclear transporters in the interactome, pointing to disruptions in nuclear transport processes in the presence of polyQ-ataxin-1. Our direct assessments of nuclear transporters and their cargoes reinforce these observations, revealing disrupted trafficking often with relocalisation of transporters and/or cargoes to ataxin-1[85Q] nuclear bodies. Strikingly, the nucleoporin Nup98, dependent on its GLFG repeats, is recruited into polyQ-ataxin-1 nuclear bodies. Our results highlight a disruption of multiple essential nuclear protein trafficking pathways by polyQataxin-1, a key contribution to furthering understanding of pathogenic mechanisms initiated by polyQ tract proteins.

## Introduction

Spinocerebellar ataxia type 1 (SCA1) is an autosomal dominant neurodegenerative disease, associated with disabilities in coordination and movement and a marked atrophy in the cerebellum and brainstem ^1^. The genetic cause underlying SCA1 has been mapped to *ATXN1*, the gene encoding the ataxin-1 protein, whereby the CAG nucleotide repeat region of *ATXN1* is expanded in SCA1 patients ^2^. The resulting polyglutamine (polyQ) tract form of the ataxin-1 protein, polyQ-ataxin-1, forms distinctive nuclear bodies in individuals with SCA1, a feature recapitulated in SCA1 transgenic mice ^3^. More broadly, polyQ tract expansions in specific proteins are now appreciated to drive at least 10 diseases ^4^, and their study is providing exciting new insights that extend beyond critical aspects of protein biochemistry including protein folding/misfolding and protein-protein interactions to important points of regulation in cellular homeostasis dictated by proteostasis and impacts on cell survival/death-decision making ^5,6,7,8,9,10,11,12^.

Several studies have reported disruptions in nuclear import/export processes in neurodegenerative diseases such as amyotrophic lateral sclerosis (ALS) and Huntington’s disease (HD) ^13,14,15,16^. The nuclear transport machinery responsible for the regulated trafficking of proteins between the cytoplasm and the nucleus has a number of key components: nuclear import/export signals (NLS/NES) of the cargo proteins that direct their nuclear/cytoplasmic distributions, dedicated transport proteins responsible for nuclear import (importins) and export (exportins), the nuclear pore complex that spans the nuclear envelope and provides a regulated gateway for nuclear trafficking events, and the RanGTP/RanGDP system that drives directionality of the transport events ^17,18^. Disrupting any of these entities can influence nucleocytoplasmic trafficking ^19,20,21^, making each of these a potential player in altered nuclear trafficking in neurodegenerative disease.

Here we approach the issues of the actions of polyQ-ataxin-1 from the perspective of protein-interaction networks, by first defining the interactome of ataxin-1[85Q]. By combining direct interaction analyses with our identification of polyQ-ataxin-1 proximal partners and pathways analyses, we not only confirm several known ataxin-1 interacting partners but we identify the nuclear transport pathway as the top-ranked cellular process defined by these interactors. By sequestering additional proteins into ataxin-1 nuclear bodies, polyQ-ataxin-1 has the potential to disrupt cellular homeostasis ^22,23,24^. To assess this possibility of ataxin-1 driven nuclear transport disruption, we define the relocalisation of multiple components of the nuclear transport machinery, often with their redistribution to ataxin-1[85Q] nuclear bodies. Our results reinforce a direct sequestration model for the impact of polyQ-ataxin-1 nuclear bodies.

## Results and Discussion

### Ataxin-1[85Q] forms distinctive nuclear bodies that are enhanced by arsenite stress

To define the suitability of the polyQ-ataxin-1 constructs GFP-ataxin-1[85Q] and MBI-ataxin-1[85Q] for interactome analyses in Neuro-2a cells, we assessed the subcellular localization of these proteins, compared to GFP or MBI alone, under control conditions and further in response to the pro-oxidant stressor arsenite ^25,26^. Whilst GFP remains broadly distributed throughout the cell, GFP-ataxin-1[85Q] forms distinctive nuclear bodies that have been reported across a range of cell types ^22,27,28^; these GFP-ataxin-1[85Q] nuclear bodies formed within 24 h of transfection and increased in size upon acute arsenite exposure (300 μM, 1 h) further highlighting their responsiveness to altered environmental conditions (Figure 1A).

**Figure 1.**
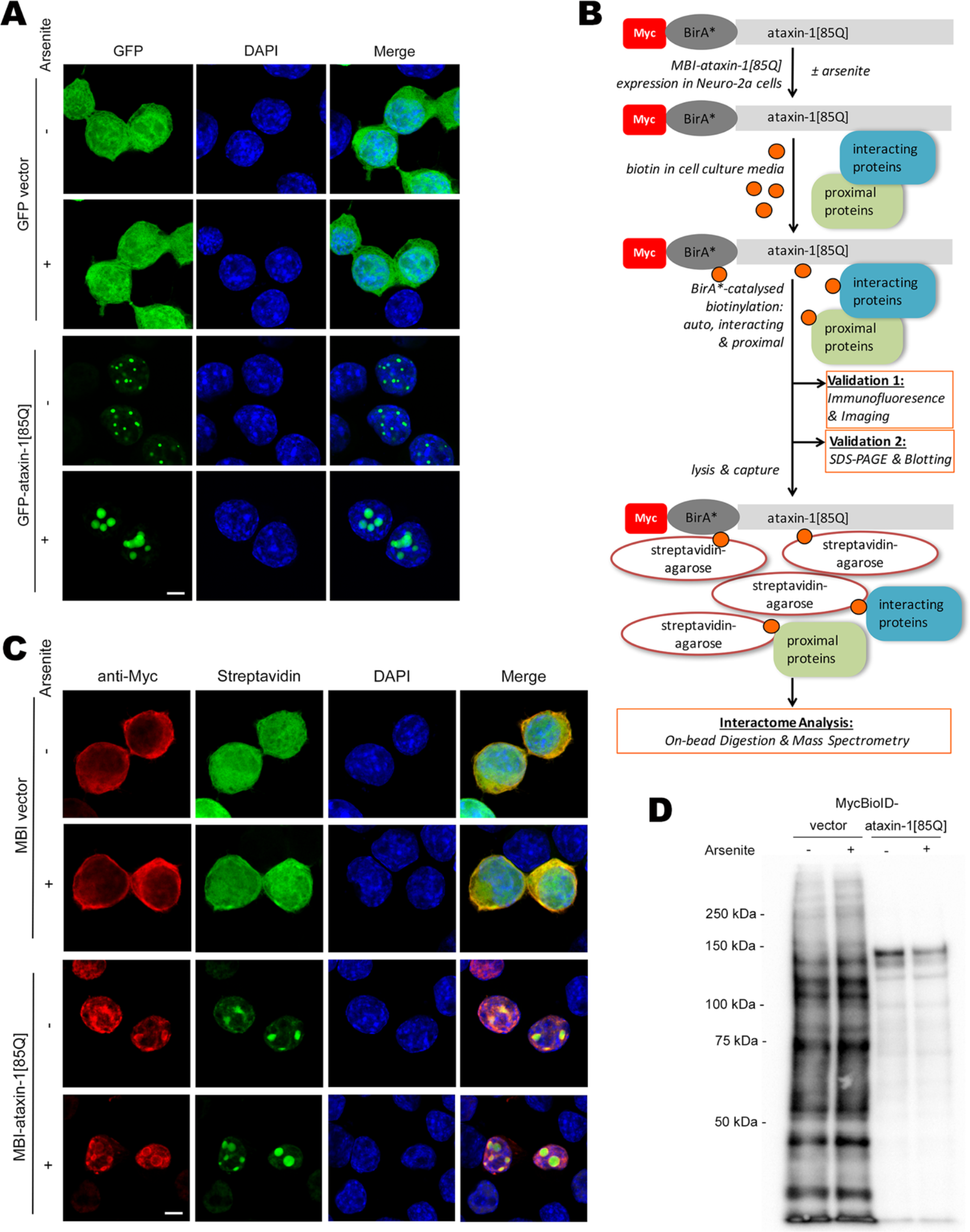
Ataxin-1[85Q] forms distinctive nuclear bodies enhanced by arsenite-induced stress. Neuro-2a cells were transfected to express either GFP-vector or GFP-ataxin-1[85Q] (A) or MBI-vector or MBI-ataxin-1[85Q] with simultaneous addition of biotin (50 μM) (B-D). At 24 h post-transfection, cells were treated with arsenite (300 μM, 1 h) as indicated. Cells were then fixed and stained with DAPI (A), or fixed, stained with anti-myc antibody, Alexa 488-streptavidin and DAPI (C), before confocal laser scanning microscopy (CLSM) imaging. Representative images are shown from 3 independent experiments; merge panels overlay GFP and DAPI images (A) or anti-myc, streptavidin and DAPI images (C), respectively. Scale bars = 10 μm. (B) Workflow for BioID sample validation and ataxin-1[85Q] interactome analysis in Neuro-2a cells. (D) At 24 h post-transfection, cells were lysed and lysates subjected to SDS-PAGE, with biotinylated proteins subsequently detected using streptavidin-HRP. Results are typical of 2 independent experiments.

Proximity biotinylation as driven by BirA* biotin ligase provides an alternative proteomics strategy for proximal partner identification ^29^. MBI-ataxin-1[85Q], the myc-tagged BirA* N-terminal fusion with ataxin-1[85Q], was thus validated as per our workflow (Figure 1B). In contrast to MBI alone that was primarily cytoplasmic, MBI-ataxin-1[85Q] was largely restricted to the nucleus (Figure 1C). Streptavidin-based detection further confirmed widespread biotinylation by MBI alone whereas MBI-ataxin-1[85Q]-driven biotinylation was largely localised to prominent nuclear bodies (Figure 1C) with a restricted biotinylation profile as defined by streptavidin-detection of biotinylated proteins separated by SDS-PAGE (Figure 1D). Thus, we confirmed the formation of distinctive nuclear bodies by GFP-ataxin-1[85Q] or MBI-ataxin-1[85Q] and the suitability of these proteins for more detailed analyses of the polyQ-ataxin-1 interactome.

### Complementary proteomics approaches identify nuclear transport proteins in the polyQ-ataxin-1 interactome

In subsequent analyses, we combined the power of BioID and Pulldown protocols as alternative and complementary approaches for interactome analyses ^30^ using both MBI-ataxin-1[85Q] and GFP-ataxin-1[85Q] in Neuro-2a cells under control and arsenite stress conditions (Figure 2A). Following trypsin-digestion of the proteins captured in the two different isolation protocols, our mass spectrometry and peptide raw data analysis defined the proteins present in each individual sample (Figure 2B). From the samples analysed as biological triplicates across the 4 conditions (BioID ± arsenite, Pulldown ± arsenite), and with the removal of non-specific proteins identified (see Methods), we recorded a total of 675 proteins (Supplementary Table S1), extending the breadth of the interactome of polyQ-ataxin-1, as previously investigated in non-neuronal (HEK293T) cell lysates co-immunoprecipitated with myc-ataxin-1 ^31^. Of note, we identified ataxin-1 itself, as well the well-characterised ataxin-1 partner protein Capicua transcriptional repressor (CIC) ^5,32,33,34^ under all 4 conditions, supporting the robustness of our approaches.

**Figure 2.**
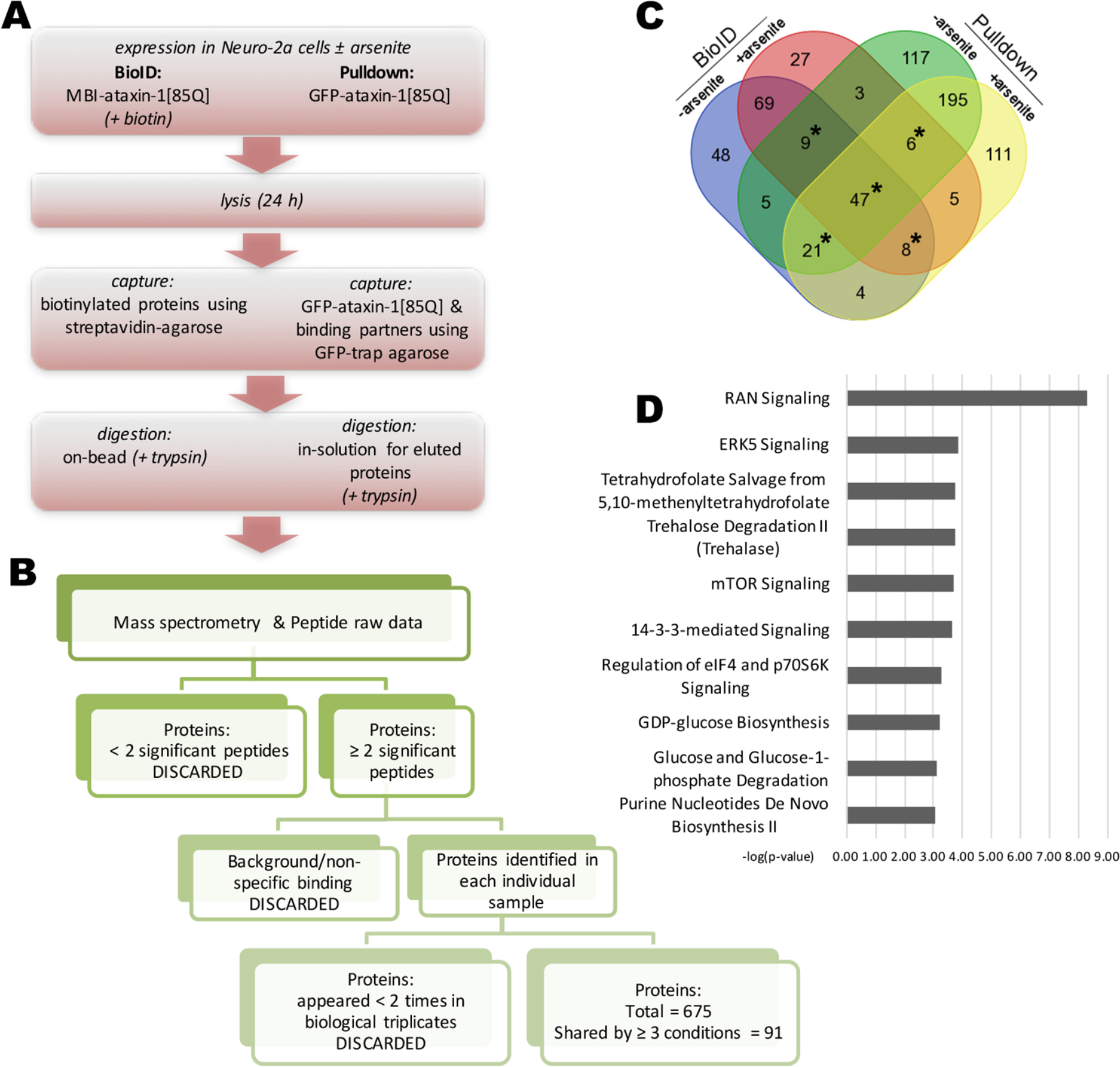
Combined approaches to ataxin-1[85Q] interactome identification reveal enrichment of nuclear transport proteins. (A, B) Parallel workflows for sample preparation for mass spectrometry to identify the ataxin-1[85Q] interactome in Neuro-2a cells (± arsenite treatment, 300 μM, 1 h). (A) The BioID protocol included the control without transfection or the expression of MBI-ataxin-1[85Q] with the simultaneous addition of biotin (50 μM) for 24 h, whereas the Pulldown protocol included the expression of GFP or GFP-ataxin-1[85Q] for 24 h. Following cell lysis, proteins were captured then subjected to trypsin digestion as indicated. (B) Mass spectrometry MASCOT analysis identified peptides from the 4 treatment conditions (BioID ± arsenite; Pulldown ± arsenite) that were matched to the protein reference library SWISSPROT (Mus musculus). Refinement of identified proteins included discarding those proteins identified by < 2 peptides and those that appeared in background control samples (i.e. for no transfection or GFP alone). Of 675 proteins identified, 91 proteins were consistently shared by ≥ 3 conditions across biological triplicate samples. (C) Venn diagram overview of the 675 identified proteins, grouped according to cell exposure (± arsenite treatment) and sample preparation (BioID or Pulldown). Areas marked by asterisk indicate those groupings of proteins identified in ≥3 conditions. (D) Ingenuity Pathway Analysis (IPA) was performed on the 91 proteins consistently shared by ≥ 3 conditions across biological triplicate samples. The top-ranked category as assessed by the −log(p-value), RAN Signaling, represents nuclear transport proteins.

Of the 675 proteins of the ataxin-1 interactome, 91 were observed in at least 3 out of 4 of the conditions analysed (Figure 2B & C), Ingenuity Pathway Analysis revealing the top-ranked pathway to be Ran signaling (Figure 2D; see summary in Supplementary Table S2). Previous studies have reported the ALS-causing C9orf72 repeat expansion RNA product to interact directly with the Ran regulator Ran GTPase activating protein 1 (RanGAP1), resulting in its mislocalization and disruption of nucleocytoplasmic transport ^14^. Significantly, RanGAP1, as well as the NPC component nucleoporin NUP62, colocalize with huntingtin protein aggregates^16^, prompting us to examine the potential impact of polyQ-ataxin-1 on nuclear transport processes.

### Classical nuclear transport pathways are disrupted by polyQ-ataxin-1

To begin to define the impact of polyQ-ataxin-1 on nuclear transport, we first examined well-characterised members of the importin (IMP) superfamily of nuclear transporters, importin-α2 (IMPα2), importin-β1 (IMPβ1) and exportin-1 (also known as CRM1). In classical nuclear protein import, IMPβ1 mediates the transfer of cargo proteins across the nuclear pore complex either directly or in conjunction with IMPα2 that acts as the adaptor binding both the cargo protein’s NLS and IMPβ1 ^19^. When bound to RanGTP, CRM1 specifically recognises leucine-rich NESs to mediate protein export from the nucleus ^35^. We coexpressed ataxin-1[85Q] with GFP-IMPα2, -IMPβ1, -CRM1, or GFP alone as a control to assess the effect of ataxin-1[85Q] on nucleocytoplasmic distribution under either control or arsenite stress conditions (Figure 3). In the presence or absence of arsenite, GFP-IMPα2 was primarily nuclear, as previously observed ^36^, while GFP-IMPβ1 and -CRM1 were primarily cytoplasmic with a distinctive nuclear rim ^37,38^; GFP was distributed throughout the cytoplasm and nucleus. In the presence of ataxin-1[85Q], however, GFP-IMPα2, -IMPβ1 and -CRM1 all showed prominent nuclear co-localisation with ataxin-1[85Q] that was further observed as striking nuclear bodies following arsenite stress exposure (Figure 3A-C, denoted by arrowheads in Zoom panels); no pronounced nuclear body distribution was observed for GFP alone (Figure 3D).

**Figure 3.**
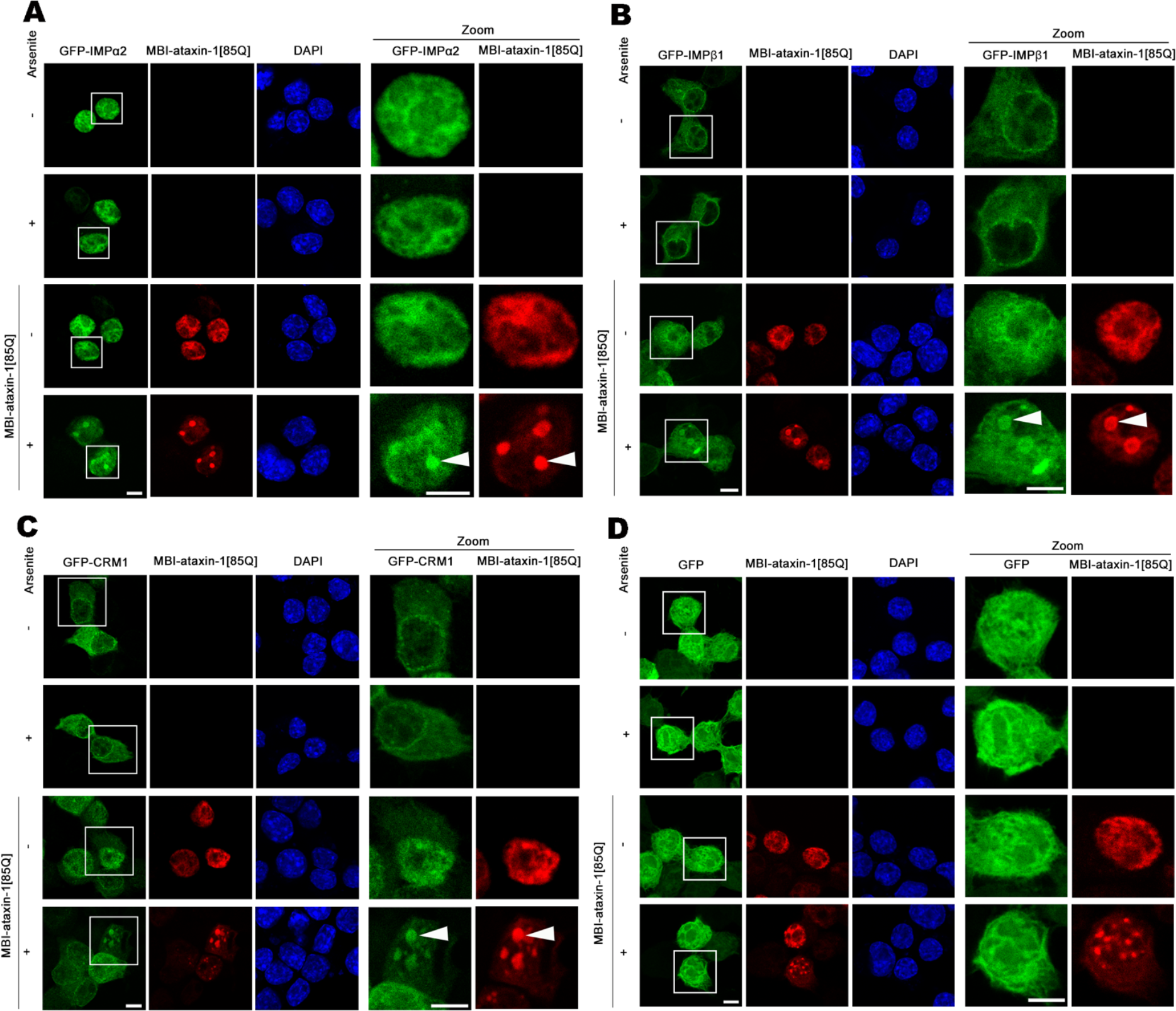
The classical nuclear import proteins, importin-α2 and importin-β1, as well as the exporter exportin-1, partially co-localise with the ataxin-1[85Q] nuclear structures. Neuro-2a cells were cotransfected to express MBI-ataxin-1[85Q] together with (A) GFP-Importin-α2 (IMPα2), (B) GFP-Importin-β1 (IMPβ1), (C) GFP-CRM1, or (D) GFP alone. At 24 h post-transfection., cells were treated with arsenite as indicated, and then fixed, stained with anti-myc antibodies and DAPI, before CLSM imaging. Representative images of cells from 2 independent experiments are shown, with zoom images (right panels) corresponding to the boxed regions. Increased co-localization, by accumulation in the ataxin-1 nuclear bodies, is indicated by the white arrowheads. Scale bar = 10 μm.

Mislocalisation of nuclear transporters would be expected to result in altered nucleocytoplasmic distribution of specific cargo proteins. Accordingly, we tested the impact of ataxin-1[85Q] expression on the nuclear accumulation of a classical importin α/β1-recognised model cargo, GFP-NLS-βgal, containing the NLS from simian virus SV40 large tumour antigen (Figure 4). We observed prominent nuclear localization of GFP-NLS-βgal under both control and arsenite stress conditions, as expected, but noted increased cytoplasmic levels of this protein under both conditions in the presence of ataxin-1[85Q] (Figure 4A, denoted by arrows). This was confirmed by image analysis to determine the nuclear to cytoplasmic ratio (Fn/c) (Figure 4B).

**Figure 4.**
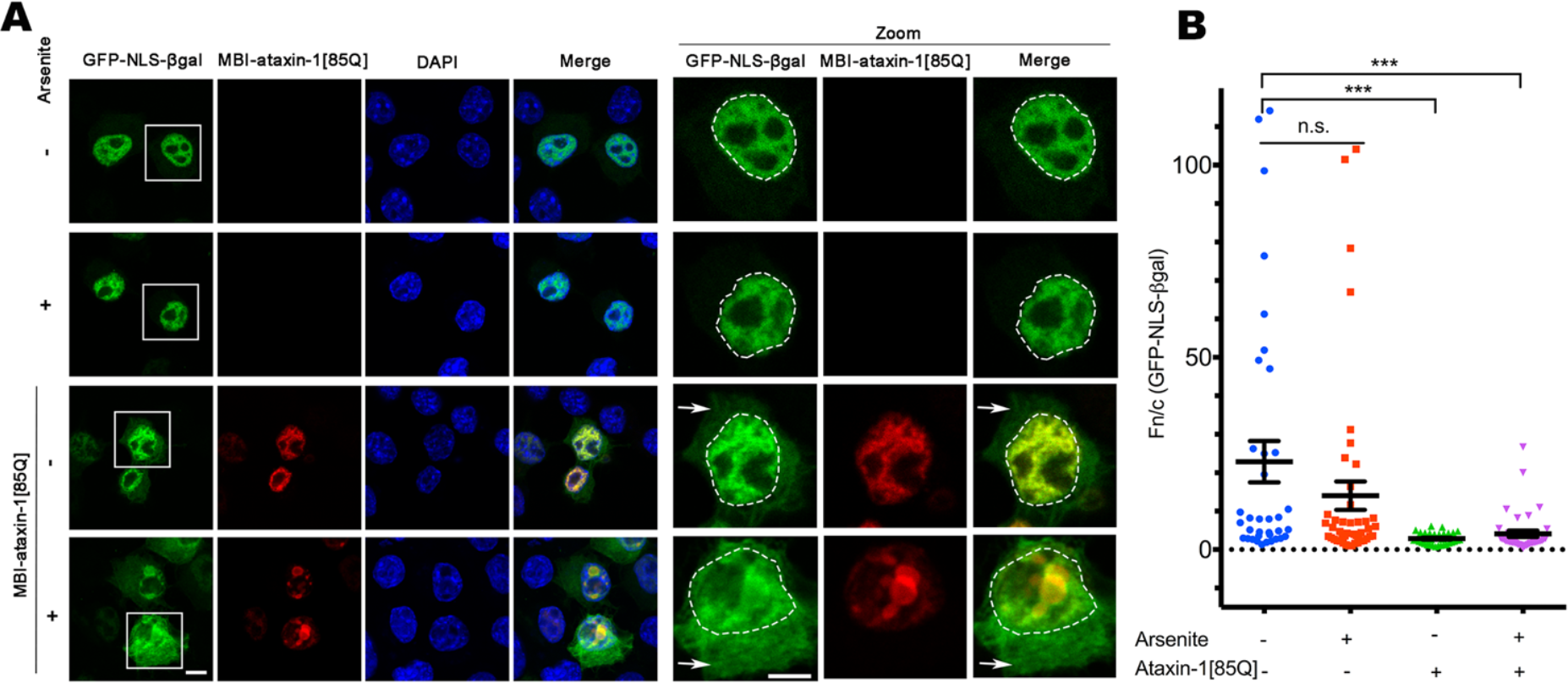
Reduced nuclear accumulation of the conventional importin-α/β1;-recognised cargo GFP-TagNLS-β-galactosidase in the presence of ataxin-1[85Q]. Neuro-2a cells were cotransfected to express MBI-ataxin-1[85Q] together with GFP-TagNLS-β-galactosidase (GFP-NLS-βGal). At 24 h post-transfection., cells were treated with arsenite as indicated, and then fixed, stained with anti-myc antibody and DAPI, before CLSM imaging. (A) Representative images from 2 independent experiments are shown; merge panels overlay GFP, myc, and DAPI images. Zoom images (right panels) correspond to the boxed regions and the position of the nucleus as determined by DAPI staining is indicated by the white dashed lines. Thin white arrows denote increased cytoplasmic fluorescence in the presence of MBI-ataxin-1[85Q]. Scale bar = 10 μm. (B) Integrated fluorescence intensity in nucleus (n) and cytoplasm (c) was estimated and the nuclear to cytoplasmic fluorescence ratio (Fn/c) determined using a modified CellProfiler pipeline. Results represent the mean ± SEM (n > 25). *** p<0.001, n.s. not significant).

### Other nuclear import pathways are selectively influenced by polyQ-ataxin-1

To extend our survey of nuclear transporters in the presence of polyQ-ataxin-1 expression at the single cell level, we next evaluated additional nuclear transport proteins. Comparable to our initial findings (Figure 3), we observed mislocalisation of both importin-13 (IMP13) ^39^ (Fig 5A), and Hikeshi, a non-importin that is responsible for nuclear import of heat shock protein 70 (HSP70) in response to heat shock ^40^ (Fig 5B). Both GFP-IMP13 and -Hikeshi were primarily nuclear in the prescence or absence of arsenite stress, but showed partial co-localization with ataxin-1[85Q] under control conditions, with colocalization with polyQ-ataxin-1 nuclear bodies in the presence of arsenite-induced stress. Consistent with these observations, we observed alterations in the nucleocytoplasmic distribution for the IMP13 cargo, transcription factor NF-YB ^41^ (Fig 5C), and the Hikeshi cargo, HSP70 ^40^ (Fig 5D), both showing partial nuclear colocalization with ataxin-1[85Q] that was enhanced by arsenite stress treatment (Figure 5C, D).

**Figure 5.**
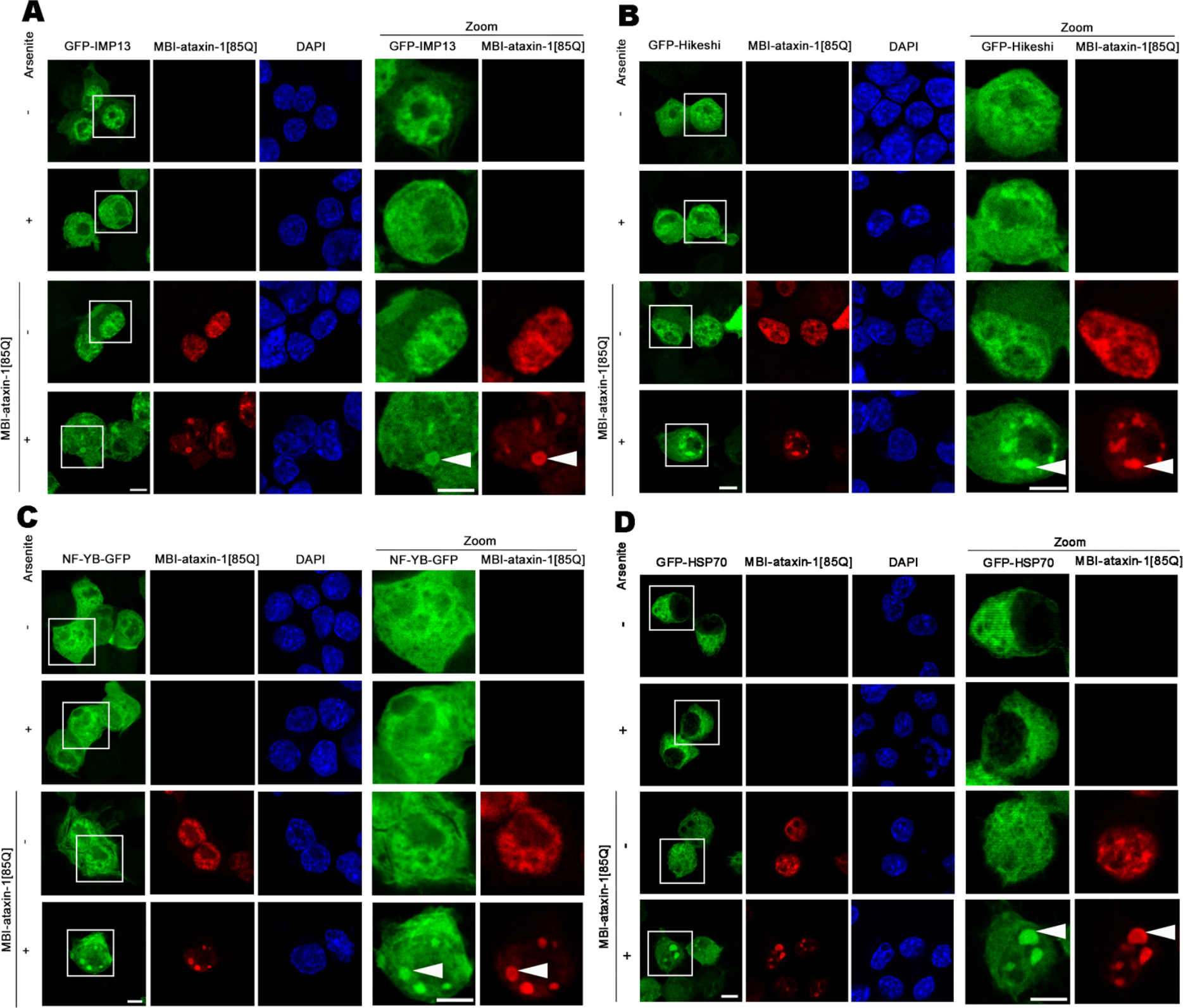
Ataxin-1[85Q] alters Importin-13 and Hikeshi localization, impacting their transport activity. Neuro-2a cells were transfected to coexpress MBI-ataxin-1[85Q] with (A) GFP-Importin-13 (GFP-IMP13), (B) GFP-Hikeshi, (C) IMP13 cargo protein NF-YB-GFP or (D) Hikeshi cargo protein GFP-HSP70. At 24 h post-transfection, cells were treated with arsenite as indicated, and then fixed, stained with anti-myc antibody and DAPI, before CLSM imaging. Representative images from 2 independent experiments are shown; merge panels overlay GFP, myc, and DAPI images. Zoom images (right panels) correspond to the boxed regions; colocalization in MBI-ataxin-1[85Q] nuclear bodies is denoted by the white arrowheads. Scale bar = 10 μm.

In contrast to the results above, we observed that a number of other nuclear transporters proteins were not impacted by/sequestered into polyQ-ataxin-1 nuclear bodies. Specifically, GFP-IMPα4, -IMP7, and -IMP16 did not show recruitment into the polyQ-ataxin-1 nuclear bodies under control or arsenite stress conditions (Figure S1A-C). GFP-IPO5 (IMP5), in contrast, appeared to localise to the polyQ-ataxin-1 nuclear structures, but only in the absence of arsenite-induced stress (Figure S1D, denoted by arrowheads in Zoom panel). Taken together, these observations highlight selectivity in the impact of the polyQ-ataxin-1 nuclear bodies by disrupting multiple, but by not all, nuclear import pathways.

The nuclear protein transport pathways mediated by importins/exportins are Ran-dependent and so we next assessed any impact of polyQ-ataxin-1 on RanGTP/GDP distribution through altering localization of its regulators, the Ran GTPase activating protein RanGAP1, and the Ran guanine nucleotide exchanger RCC1. RanGAP1 can localize with mutant huntingtin or C9orf72 polyGA aggregates in *in vivo* models of HD and ALS ^14,15^, but we observed no alteration in RanGAP1 subcellular distribution in the presence of polyQ-ataxin-1 expression (Figure S2). With our proteomics data identifying RanGAP1, as well as Ran binding protein 1 (RanBP1), in association with GFP-ataxin-1[85Q] in our pulldown analysis (Supplementary Table S1), the implication is that significant mislocalization of RanGAP1 requires longer term exposure to polyQ-ataxin-1 aggregates, as observed for huntingtin-RanGAP1 association in vivo ^15^. In contrast, our assessment of endogenous RCC1 or ectopically addressed GFP-RCC1 revealed clear colocalisation with polyQ-ataxin-1 nuclear bodies (Figure S3A,B). Thus, through sequestrating RCC1 into distinct nuclear bodies, polyQ-ataxin-1 has the potential to reduce nuclear RanGTP levels, and hence importin/exportin-dependent nuclear trafficking. Additionally, not all RCC1 is sequestered into polyQ-ataxin-1 nuclear bodies, explaining how there is not complete disruption of the localization of all nuclear transporters/nuclear transport pathways. Furthermore, that Ran-independent (Hikeshi) and Ran-dependent (importins/exportins) nuclear transport pathways are impacted by polyQ-ataxin-1 expression suggests that there may be additional effects at the level of the nuclear pore complex (NPC).

### PolyQ-ataxin-1 expression results in mislocalisation of nucleoporin NUP98

The NPC is the regulated gateway for trafficking between the cell cytosol and nucleus, and is a potential target of mutant polyQ-huntingtin protein in HD ^16^, where progressive disruption of nuclear envelope morphology has been observed in aging mice expressing the polyQ-huntingtin protein ^15^. Post-mortem samples from patients with ALS ^42^ also show disruptions to the nuclear envelope, whereas polyQ-ataxin-1 may also decrease nuclear membrane instability^43^.

We set out to evaluate the effect of short-term expression of polyQ-ataxin-1, initially by assessing NPC distribution using the anti-nucleoporin (Nup) Mab414 antibody; no marked changes in staining were observed upon polyQ-ataxin-1 expression (Figure 6A) indicating no wide-spread disruption of NPC distribution in the nuclear envelope. Indicative of general NPC functionality, nuclear accumulation of the transcription factor AF10, known to be importin/Ran-independent and to occur via direct interaction with NPC components ^44,45^ was not affected by polyQ-ataxin-1 (Figure S4). Nucleoporin protein NUP62, within the central core of the NPC ^46,47^, was not altered in the presence of polyQ-ataxin-1 either in the presence or absence of arsenite stress (Figure S5), in contrast to the impact of mutant huntingtin in HD^16^. Importantly, although NUP62 staining showed distinctive nuclear puncta, these did not co-localise with the polyQ-ataxin-1 nuclear bodies (Figure S5), indicating that the effects of polyQ-ataxin-1 and mutant huntingtin are distinct, although both impact nuclear transport machinery.

**Figure 6.**
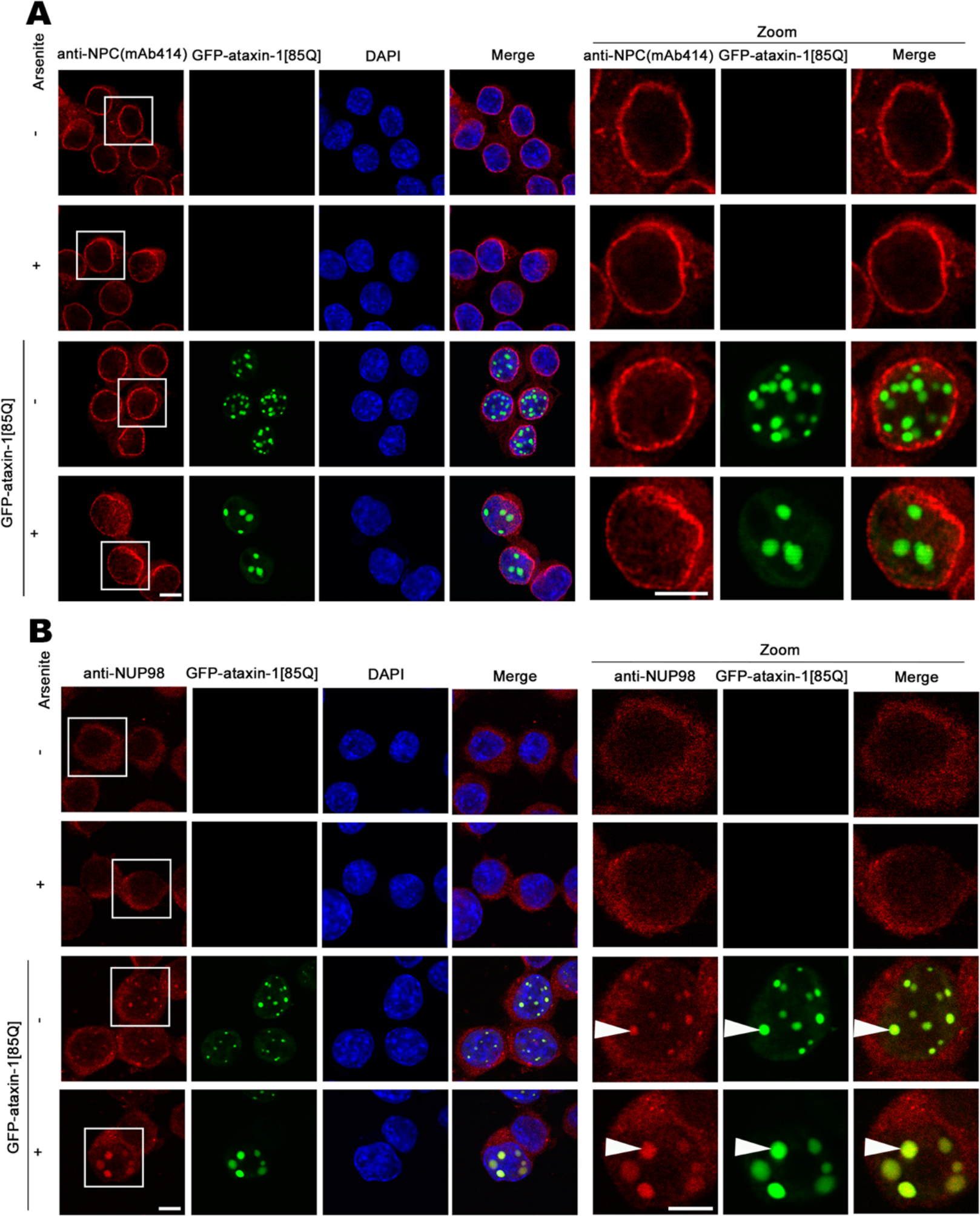
Ataxin-1[85Q] expression alters NUP98 localization but not general NPC distribution. Neuro-2a cells were transfected to express MBI-ataxin-1[85Q] and treated with arsenite as indicated at 24 h post-transfection. Cells were then fixed, stained with anti-NPC (Mab414) (A) or anti-NUP98 (B) antibodies together with DAPI, before CLSM imaging. Representative images from 2 independent experiments are shown; merge panels overlay (A) NPC, GFP, and DAPI or (B) NUP98, GFP, and DAPI images. Zoom images (right panels) correspond to the boxed regions; white arrowheads denote colocalisation in ataxin-1[85Q] nuclear bodies. Scale bar = 10 μm.

We next evaluated NUP98, the most conserved FG-dipeptide motif domain nucleoporin and a major contributor to the NPC permeability barrier ^48,49^. Indeed, the more cohesive nature of NUP98 GLFG repeat domains compared to the FxFG repeats of other nucleoporins ^50^ can drive the formation of highly concentrated gels ^51^. Immunostaining of endogenous NUP98 revealed its distribution throughout the cell with recruitment into polyQ-ataxin-1 nuclear bodies (Figure 6B, denoted by white arrowheads in Zoom panels). Analogously, ectopically expressed GFP-NUP98, colocalised with polyQ-ataxin-1 nuclear bodies upon arsenite exposure (Figure 7A). To further dissect the mechanisms contributing to this co-localisation, we evaluated the localisation patterns of three truncation derivatives of NUP98: GFP-NUP98 N-terminal (1-225), GFP-NUP98 GLFG domain (221-504) and GFP-NUP98 C-terminal (506-920). We observed overlap but no enrichment in fluorescence colocalised with the polyQ-ataxin-1 nuclear bodies for the N-terminal or C-terminal constructs (Figure 7B). In contrast, we observed small nuclear puncta for GFP-NUP98 GLFG domain (221-504) even in the absence of polyQ-ataxin-1 (Figure 7B, denoted by open arrowheads in Zoom panels); upon arsenite treatment, striking colocalisation in the larger polyQ-ataxin-1 nuclear bodies was also evident (Figure 7B, denoted by white arrowheads in Zoom panels). Thus, sequestration of NUP98 into the polyQ-ataxin-1 nuclear bodies is dependent on the NUP98 GLFG domain.

**Figure 7.**
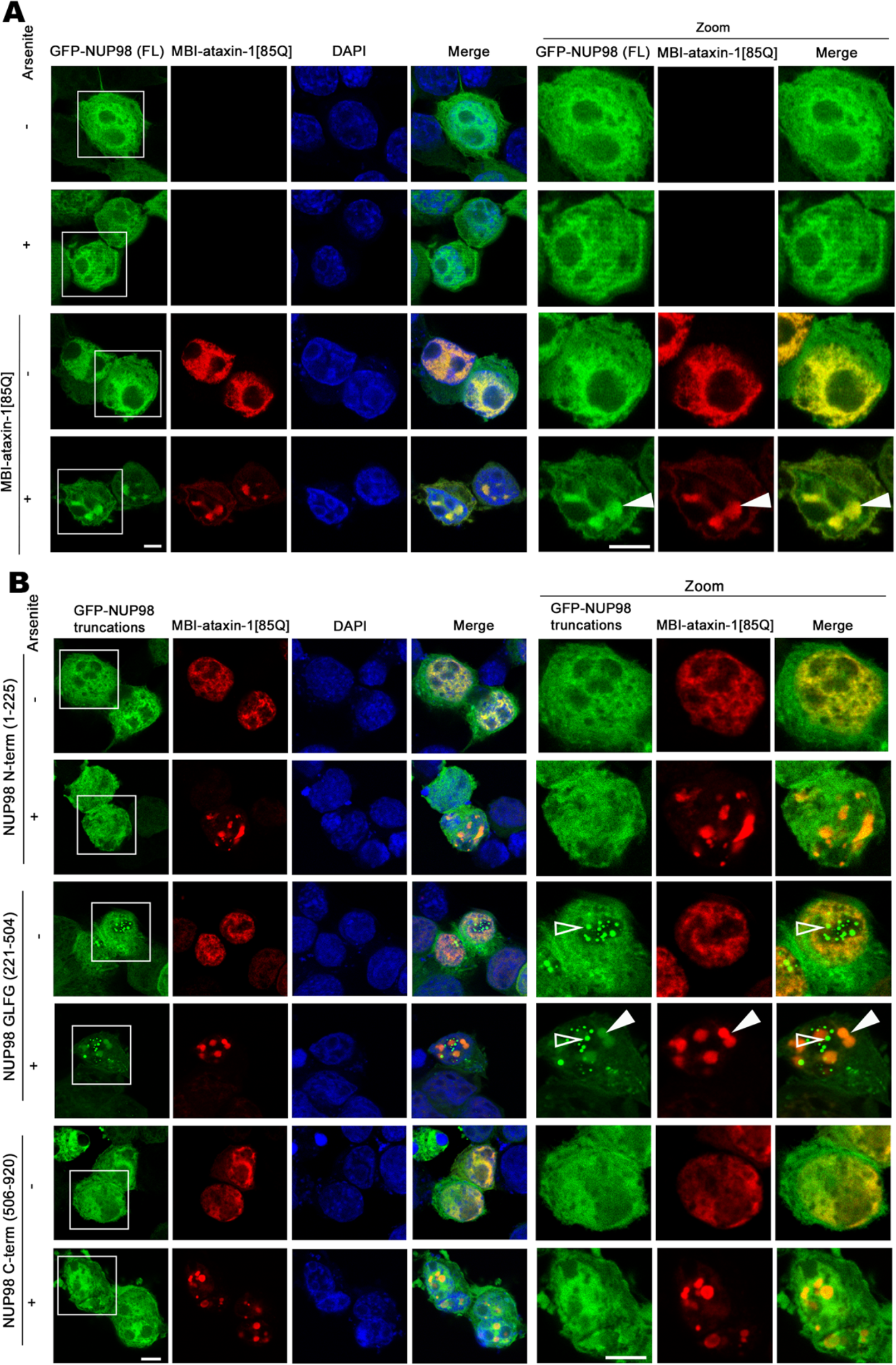
The nucleoporin NUP98 is recruited into ataxin-1[85Q] structures, and this recruitment depends on the NUP98 GLFG repeats. Neuro-2a cells were transfected to coexpress MBI-ataxin-1[85Q] with GFP-NUP98 (A), or the indicated GFP-NUP98 truncation constructs (B). At 24 h post-transfection., cells were treated with arsenite as indicated, and then fixed, stained with anti-myc-antibodies and DAPI, before CLSM imaging. Representative images from 2 independent experiments are shown; merge panels overlay GFP, myc, and DAPI images. Zoom images (right panels) correspond to the boxed regions. White arrowheads denote colocalization in ataxin-1[85Q] nuclear bodies; NUP98 nuclear puncta are indicated by the open arrowheads. Scale bar = 10 μm.

Previous studies using knockdown of NUP98 to interrogate its functions have shown selective disruption of Tag-NLS-/importin α/β1-dependent and M9-NLS-/transportin-1 (TNPO1)-dependent nuclear protein import, but not BIB (“β-importin-binding”)-NLS-dependent transport, that can be mediated by multiple different importin βs ^20,52^. To test the possible impact of NUP98 sequestration by polyQ-ataxin-1 nuclear bodies on nuclear transport, we examined the subcellular localisation of GFP-M9-NLS and GFP-BIB-NLS. We found that the extent of nuclear accumulation of GFP-M9-NLS, but not of GFP-BIB-NLS was significantly (p < 0.05) increased in the presence of polyQ-ataxin-1 expression and arsenite induced stress (Figure S6). These results further support the observations that some, but not all, nuclear transport processes are disrupted by polyQ-ataxin-1.

### The ataxin-1 interactome indicates impacts on multiple intracellular processes beyond nuclear transport

Previous genetic screens have highlighted the impact of nuclear transport proteins across a number of neurodegenerative diseases. In exploring suppressors of C9orf72 repeat expansion toxicity in *Drosophila* ALS model, multiple nuclear pore complex proteins, importins/exportins and Ran regulators have been identified ^13,14,53^. The earlier screens for genetic suppressors of ataxin-1-induced neuronal degeneration in a *Drosophila* SCA1 model previously identified the nucleoporin NUP44A as a suppressor of toxicity ^54,55^. Our study is the first to investigate the impact of short-term polyQ-ataxin-1 expression on cellular nuclear transport processes, our findings indicating that nuclear transport is directly disrupted by short-term polyQ-ataxin-1 expression. Our proteomics analyses, in combination with our surveying of additional nuclear transporters, Ran regulators and nucleoporins, point to the disruption at multiple critical points across the cellular nuclear transport system.

Our results emphasize that approaches to rescue the nuclear transport disruption by mutant ataxin-1 may ultimately provide new ideas on the possible therapeutic interventions to slow neurodegenerative disease progression in SCA1. Furthermore, the ataxin-1 interactome remains a resource for further exploration. For example, we identified replication protein A1 (Rpa1) by pulldown approaches (Table S1); Rpa1 has been previously identified as having the largest effect on lifespan in the *Drosophila* SCA1 model and has been located at the hub position linked to repair systems such as homologous recombination ^56^. Others have implicated genes in the protein folding/heat-shock response or ubiquitin-proteolytic pathways as modifiers of toxicity in models of SCA1 ^54^ and our proteomics studies have identified multiple heat shock proteins as well as proteins involved in ubiquitination, and ubiquitin itself, as part of the ataxin-1 interactome (Table S1). Ataxin-1 phosphorylation may also influence ataxin-1 stability ^57^; with the implication of cAMP-dependent protein kinase as a mediator of phosphorylation and toxicity of ataxin-1 ^58,59^, it is of interest that we identified both regulatory and catalytic subunits of cAMP-dependent protein kinase in our interactome analysis (Supplementary Table S1). Increasing levels of wild-type ataxin-1 are sufficient to lead to SCA1-like neurodegeneration^60^, and thus the exact balance of binding partner selection by ataxin-1 is of critical importance. Deciphering the cellular events underlying the ataxin-1 interactome may well be the key to new treatment options for SCA1 and other neurodegenerative diseases.

## Materials and methods

### Plasmids

The GFP-ataxin-1[85Q] plasmid was provided by D Hatters (University of Melbourne). Myc-BioID (MBI)-ataxin-1[85Q], the myc-tagged BirA* N-terminal fusion with ataxin-1[85Q], was generated by amplifying the coding region of ataxin-1[85Q] from the GFP-ataxin-1[85Q] construct to include the desired restriction enzyme sites (EcoRI and HindIII), and then ligation into the pcDNA3.1_mycBioID plasmid ^29^ (#35700, addgene). The PCR primer pairs used were 5’-GCGAATTCATGAAATCCAACCAAGAGCGGAGC-3’ (forward) and 5’- GCAAGCTTCTACTTGCCTACATTAGACCGGCC-3’ (reverse). All constructs were validated by restriction digestion and full sequence analysis.

### Cell culture, transfection and treatment

Mouse neuroblastoma cells (Neuro-2a) were cultured in Opti-MEM supplemented with 2 mM L-glutamine (Gibco), 10% fetal bovine serum (BOVOGEN) and 100 U/ml penicillin/streptomycin (Gibco). Cells were plated and cultured (16 h), transfected with the indicated constructs using Lipofectamine 2000 reagent (Invitrogen) according to the manufacturer’s instructions (24 h), and then exposed as indicated to 300μM sodium arsenite (Sigma) for 1 h. GFP was included as a control for imaging and proteomics protocols examining GFP-ataxin-1[85Q]. When BioID protocols were used, biotin (50 μm) was included during the transfection protocols with the MBI vector (for blot detection) or MBI-ataxin-1[85Q] (for blot detection or proteomics protocols).

### Cell lysate preparation and pull-down protocols

Cell lysates were prepared using RIPA buffer [50 mM Tris-HCl, pH 7.3, 150 mM NaCl, 0.1 mM ethylenediaminetetraacetic acid (EDTA), 1% sodium deoxycholate, 1% Triton X-100, 0.2% NaF and 100 µM Na_3_VO_4_] supplemented with complete protease inhibitor mix (Roche Diagnostic). Cell lysates were incubated on ice (20 min) and cleared by centrifugation (1,000 rpm, 20 min). Protein concentrations were determined using the BioRad assay.

For BioID pull-down prior to mass spectrometry or immunoblot analysis, whole cell lysates (3 mg protein) prepared from Neuro-2a cells expressing MBI-ataxin-1[85Q] were incubated with streptavidin-agarose (0.05 mL) (Invitrogen). For GFP pull-down prior to mass spectrometry or immunoblot analysis, whole cell lysates prepared from Neuro-2a cells expressing GFP fusion constructs (3 mg protein) were incubated with GFP-trap-agarose (0.05 mL) (ChromoTek). The incubation for streptavidin-agarose was overnight at 4ºC, and for GFP-trap-agarose was 2 h at room temperature. The immobilized proteins in the streptavidin- or GFP-trap-agarose pellets were thoroughly washed. For both protocols, 7 wash steps were performed, with centrifugation (12,000 rpm, 1 min) between washes and with the first 2 washes using RIPA buffer. For the BioID pull-down, the 5 subsequent wash steps included: 0.5% SDS in PBS (washes 3 and 4), 6 M urea in 100 mM TrisHCl pH 8.5 (washes 5 and 6), and 100 mM TrisHCl pH 8.5 (wash 7). For the GFP pull-down, all 5 subsequent washes used 10 mM Tris-HCl pH 7.5, 150 mM NaCl, 0.5 mM EDTA (washes 3 to 7).

### Blot detection of biotinylated proteins

Proteins binding to the streptavidin-agarose were eluted with concentrated SDS-sample buffer (180 mM TrisHCl pH 6.8, 30% glycerol, 6% SDS, 0.06% bromophenol blue, 15% β-mercaptoethanol). Samples were boiled (5 min, 95ºC). Eluted proteins were separated by SDS-PAGE (8% polyacrylamide gels, 1.5 h) and transferred to polyvinylidene difluoride (PVDF) membranes (Amersham Life Science; 2 h, room temperature). Subsequent steps were performed by blocking with 1% bovine serum albumin (BSA) in phosphate-buffered saline (PBS; 0.5 h, room temperature), and incubating with streptavidin-HRP (Invitrogen, 1:2000; 1h, room temperature) followed by thorough washing with Tris-buffered saline Tween-20 (TBST). The membrane was finally blocked with buffer (1% BSA, 1% Triton X-100 in PBS; 5 min), washed (TBST, 3 × 1 min), and visualised using an enhanced chemiluminescence detection system (ThermoFisher Scientific). Images were captured using ChemiDoc imager (Bio-Rad) operating in a single-channel protocol.

### Mass spectrometry (MS) analysis

Proteins in the washed streptavidin-or GFP-trap-agarose pellets were further prepared for MS analysis. For streptavidin-agarose pellets, an on-bead digestion was performed by adding 20 μl trypsin-containing denaturing solution (50 mM urea, 5 mM tris(2-carboxyethyl) phosphine (TCEP) and 0.25 μg trypsin (Sigma) in 50 mM triethylammonium bicarbonate (TEAB)). After incubation (overnight, 37C, end-over-end rotation), supernatants were collected. For the GFP-trap-agarose pellets, an in-solution digestion for the eluted proteins was performed by adding the elution solution (50% aqueous 2,2,2-Trifluoroethanol (TFE)/0.05% formic acid, pH 2.0, 1mM TCEP). After incubation (5 min, room temperature), supernatants were collected and then 40 μl trypsin solution (0.25 μg trypsin, 200 mM TEAB) added before incubation (overnight, 37C, end-over-end rotation).

For each trypsin-digested sample, supernatant (10 μl) was collected and tryptic peptides were analysed by liquid chromatography-MS/MS (LC-MS/MS) using an Q-Exactive plus mass spectrometer (Thermo Scientific) fitted with nanoflow reversed-phase-HPLC (Ultimate 3000 RSLC, Dionex). The nano-LC system was equipped with an Acclaim Pepmap nano-trap column (Dionex – C18, 100 Å, 75 µm × 2 cm) and an Acclaim Pepmap RSLC analytical column (Dionex – C18, 100 Å, 75 μm × 50 cm). Typically for each LC-MS/MS experiment, 1 μL supernatant was loaded onto the enrichment (trap) column (isocratic flow 5 μl/min, 3% CH_3_CN containing 0.1% formic acid, 6 min) before the enrichment column was switched in-line with the analytical column. For LC, the eluents used were 0.1% formic acid (solvent A) and 100% CH_3_CN/0.1% formic acid (solvent B) with the following sequence of gradients: 3 to 20% B (in 95 min), 20 to 40% B (in 10 min), 40 to 80% B (in 5 min), 80% B (maintained for the final 5 min) before equilibration in 3% B prior to the next analysis (10 min). All spectra were acquired in positive mode with full scan MS1 scanning from m/z 375-1400 (70000 resolution, AGC target 3e6, maximum accumulation time 50 ms, lockmass 445.120024). The 15 most intense peptide ions with charge states ≥2-5 were isolated (isolation window 1.2 m/z) and fragmented with normalized collision energy of 30 (35000 resolution, AGC target 1e5, maximum accumulation time 50 ms). An underfill threshold was set to 2% for triggering of precursor for MS2. Dynamic exclusion was activated for 30s.

### Bioinformatic analysis

Resultant MS/MS data was analyzed using the Mascot search engine (Matrix Science version 2.4) against the SWISSPROT database (with the settings as follows: taxonomy - Mus., enzyme - Trypsin, Protein Mass - ± 20 ppm, Fragment Mass Tolerance - ± 0.2 Pa, Max Missed Cleavages: 2). Identifications in all samples (test or background/non-specific binding) were accepted for proteins with at least two significant peptides (p<0.05). Background/non-specific binding proteins were identified as follows: for the Pulldown protocol assessments, proteins identified in samples prepared from GFP only-transfected cells processed in parallel in each of the 3 replicates were defined as background/non-specific binding; for BioID assessments, proteins identified in samples prepared from untransfected cells across all 3 replicates were pooled and defined as background as endogenously biotinylated proteins. Proteins in the resulting lists were retained for further consideration when they were identified in at least three of the four tested conditions (BioID ± arsenite; Pulldown ± arsenite) were further considered. Ingenuity pathway analysis (IPA; QIAGEN) was carried out according to the supplier’s instructions with default settings.

### Immunofluorescence, confocal laser scanning microscopy and image analysis

Neuro-2a cells were cultured on coverslips (Proscitech), transfected and treated with 300 μm arsenite, as indicated. Subsequent processing (washing, permeabilization (0.2% (v/v) Triton X-100), fixation (4% (w/v) paraformaldehyde and blocking (1% (w/v) bovine serum albumin (BSA)) was performed at room temperature in phosphate-buffered saline. Cells were then incubated sequentially with primary antibodies then fluorophore-conjugated secondary antibodies or streptavidin as indicated, each in 1% (w/v) BSA in PBS for 1 h at room temperature. The primary antibodies used were as follows: anti-cMyc (sc-40, Santa Cruz); anti-NUP, Mab414 (ab24609, abcam); anti-NUP98 (sc-14153, Santa Cruz); anti-NUP62 (610498, BD Biosciences); anti-RanGAP1 (ab2081, ancam); anti-RCC1 (sc-1162, Santa Cruz). Subsequent detection used Alexa Fluor^®^ 568-conjugated secondary antibodies (A-11004, Invitrogen) or Alexa Fluor^®^ 488-conjugated Streptavidin (S11223, Invitrogen). Processed coverslips were then mounted by water-based Fluoro-Gel (Proscitech) onto glass slides for visualization by confocal laser scanning microscopy (Leica TCS SP5 with 63X 1.4 Oil Objective). Images were analyzed using Fiji software, with quantitative analysis of the fluorescence intensity of nucleus/cytoplasm ratio (Fn/c) performed using CellProfiler cell image analysis software (version 2.1.1 for Mac).

### Statistical analysis

Graphpad Prism 6 (version 6.00 for Mac) was used for statistical analysis. For analysis of >2 datasets, ANOVA was applied to compare differences among different datasets, followed by Tukey’s multiple comparisons test. Data are represented as mean ± standard error of the mean (SEM), with p <0.05 regarded as statistically significant.

## Author contributions

S.Z., D.A.J. and M.A.B conceived the work and designed the experiments. S.Z. and N.A.W. performed the experiments and all authors analyzed the data. S.Z. drafted the manuscript, and all authors contributed to discussions and reviewed the manuscript.

## Supplementary Figures

**Figure S1.**
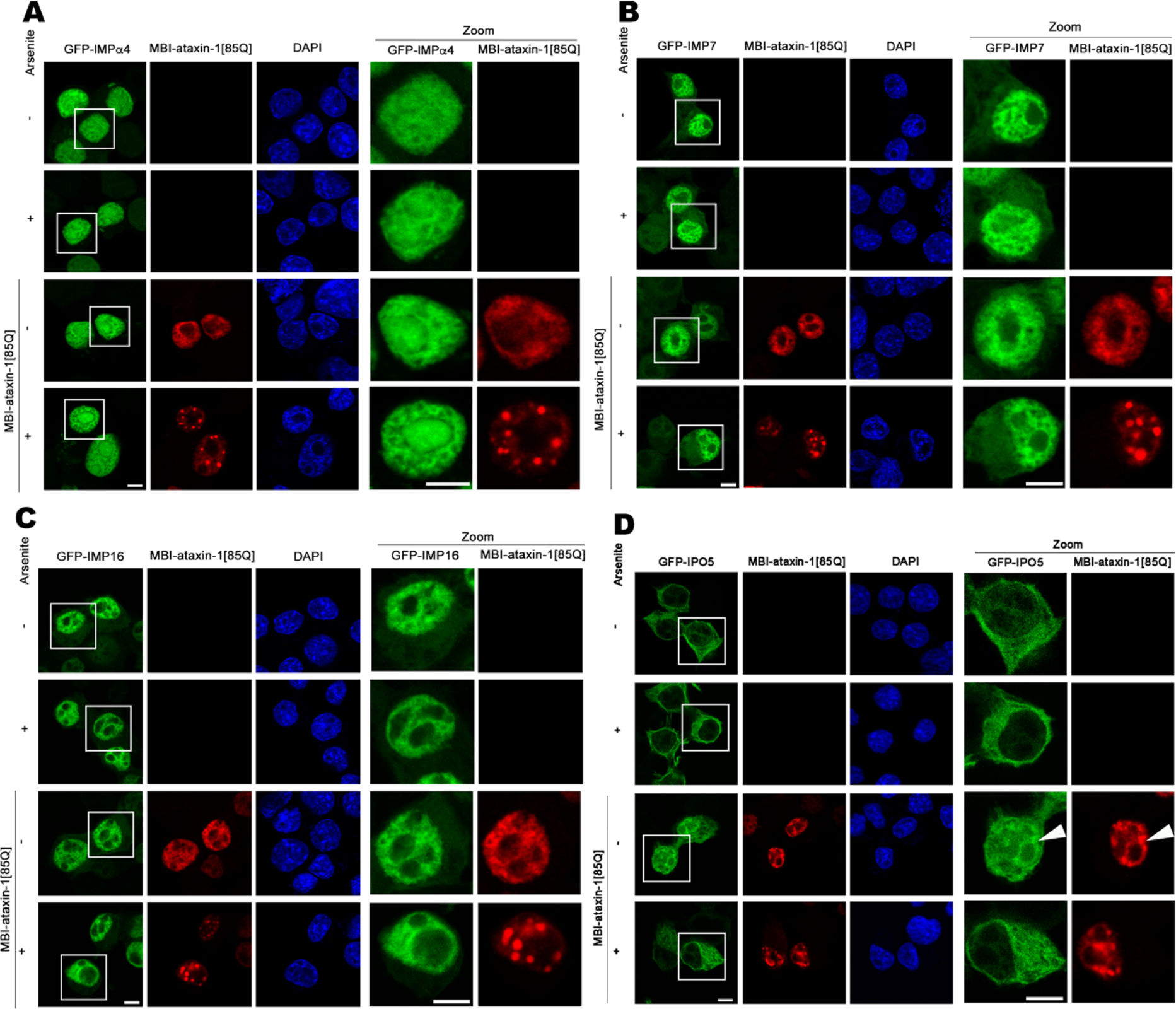
Mislocalisation of IPO5 but not IMPα4,IMP7, IMP16 upon polyQ-ataxin-1 expression. Neuro-2a cells were cotransfected to express MBI-ataxin-1[85Q] together with (A) GFP-IMPα4, (B) GFP-Importin- (IMP7), (C) GFP-Importin-16 (IMP16) or GFP-Importin-5 (IPO5). At 24 h post-transfection, cells were treated with arsenite as indicated, and then fixed, stained with anti-myc antibody and DAPI, before CLSM imaging. Representative images are shown from 2 independent experiments; zoom images (right panels) correspond to the boxed regions. Scale bar = 10 μm.

**Figure S2.**
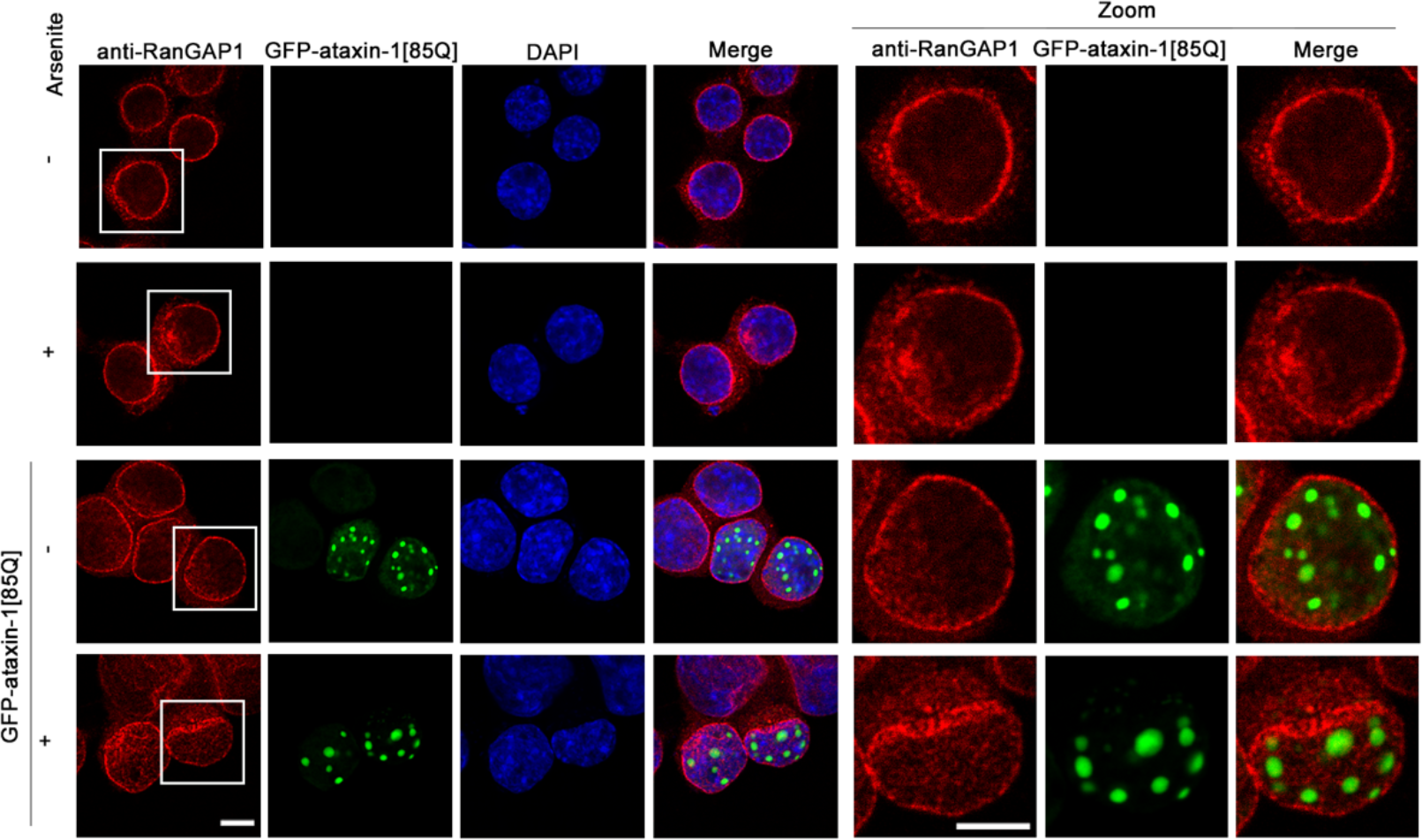
RanGAP1 localisation is unaltered upon polyQ-ataxin-1 expression. Neuro-2a cells were transfected to express GFP-ataxin-1[85Q]. At 24 h post-transfection, cells were treated with arsenite as indicated, and then fixed, stained with anti-RanGAP1 antibody and DAPI, before CLSM imaging. Representative images are shown from 2 independent experiments; merge panels overlay RanGAP1, GFP, and DAPI images. Zoom images (right panels) correspond to the boxed region. Scale bar = 10 μm.

**Figure S3.**
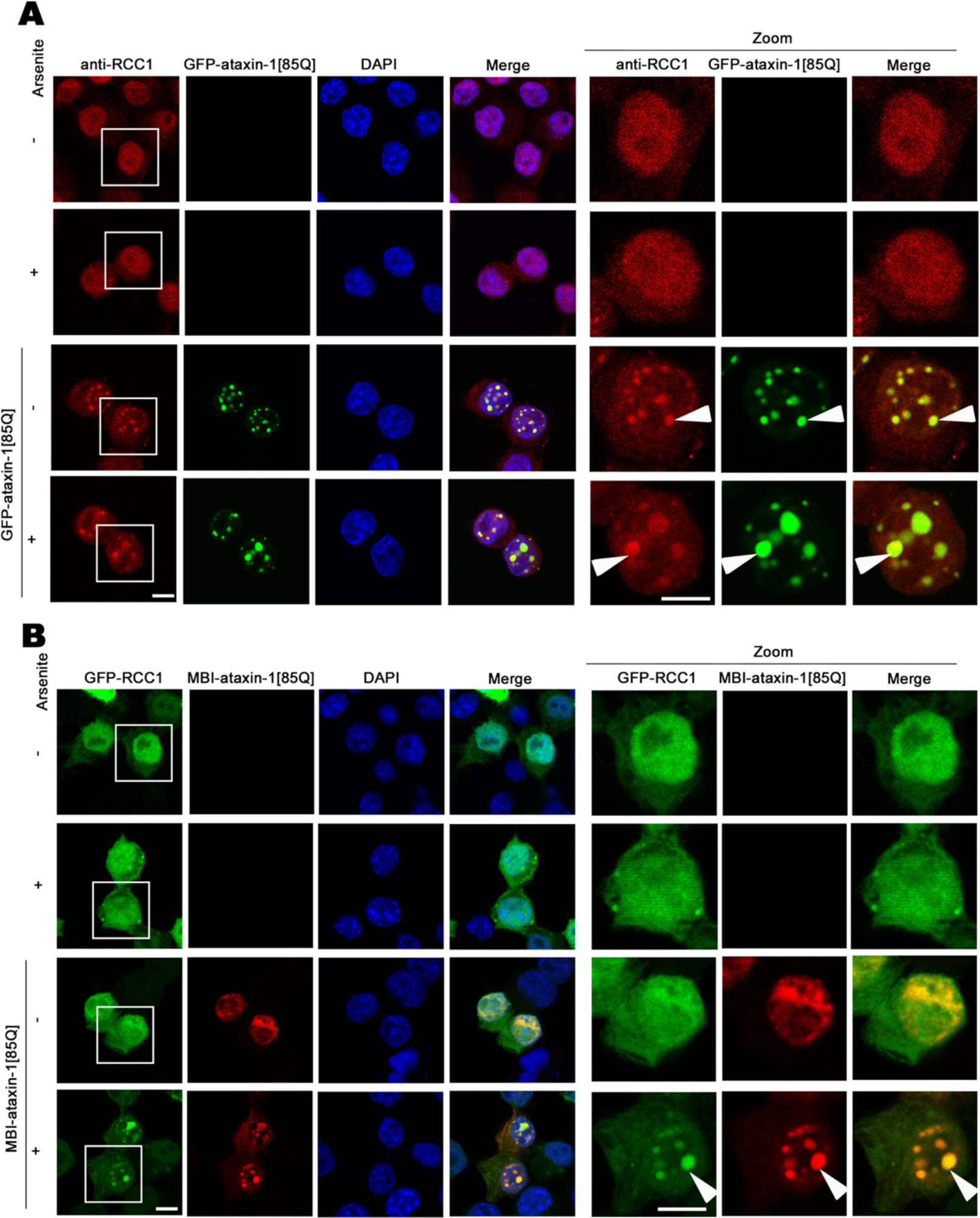
RCC1 localizes in poly-ataxin-1 nuclear bodies. Neuro-2a cells were (A) transfected to express GFP-ataxin-1[85Q] or (B) MB-ataxin-1[85Q] together with GFP-RCC1. At 24 h post-transfection, cells were treated with arsenite as indicated, and then fixed, stained with (A) anti-RCC1 or anti-myc (B) antibodies, and DAPI, before CLSM imaging. Representative images are shown from 2 independent experiments; merge panels overlay (A) RCC1, GFP, and DAPI images (B) GFP, myc, and DAPI images. Zoom images (right panels) correspond to the boxed region; colocalization in MB-ataxin-1[85Q] nuclear bodies is denoted by the white arrowheads. Scale bar = 10 μm.

**Figure S4.**
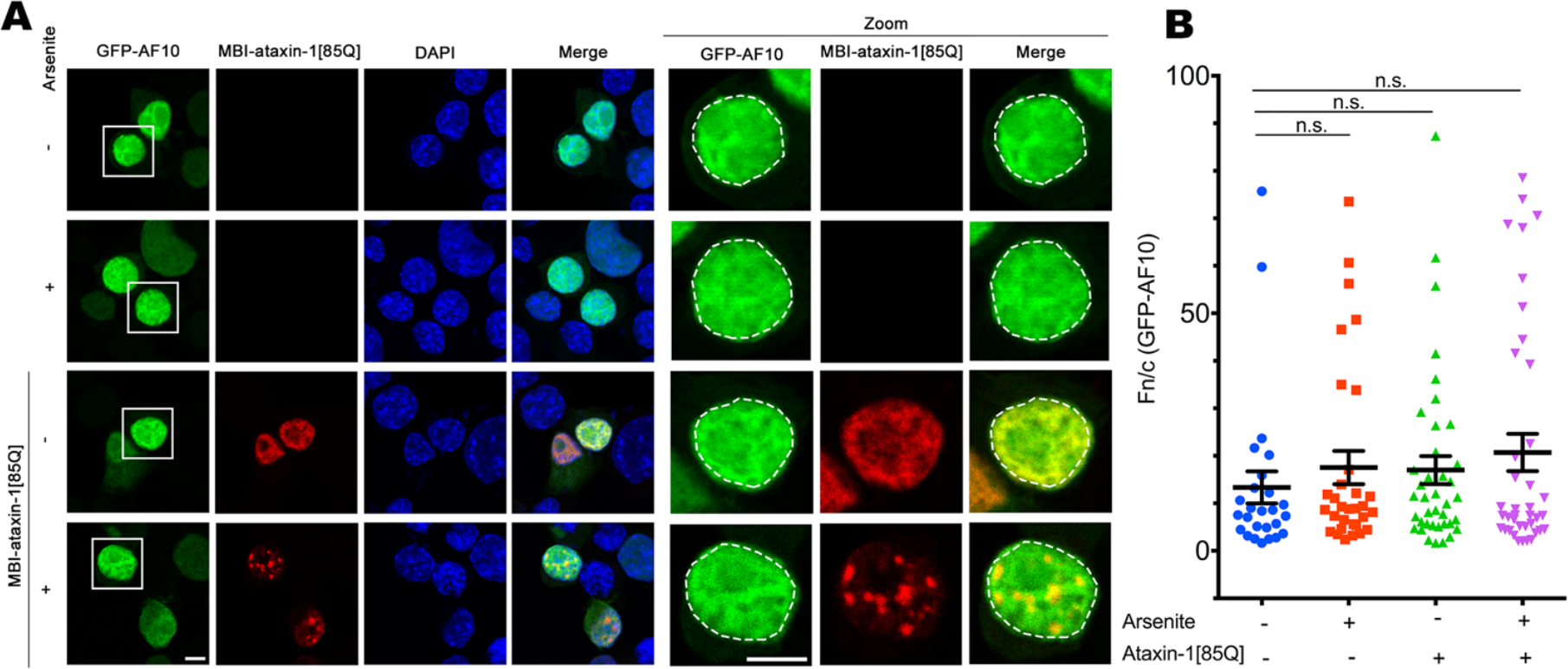
Nuclear import of GFP-AF10 is not influenced by polyQ-ataxin-1. Neuro-2a cells were transfected to coexpress MBI-ataxin-1[85Q] and GFP-AF10. At 24 h post-transfection, cells were treated with arsenite as indicated, and then fixed, stained with anti-myc-antibody and DAPI, before CLSM imaging. Representative images from 2 independent experiments are shown; merge panels overlay GFP, myc and DAPI images. Zoom images (right panels) correspond to the boxed regions. Scale bar = 10 μm. The position of the nucleus as determined by DAPI staining is indicated by the white dashed lines. The nuclear to cytoplasmic fluorescence ratio (Fn/c) was calculated by a modified CellProfiler pipeline as per the legend to Figure 4B. Results represent the mean ± SEM (n > 25); n.s. not significant.

**Figure S5.**
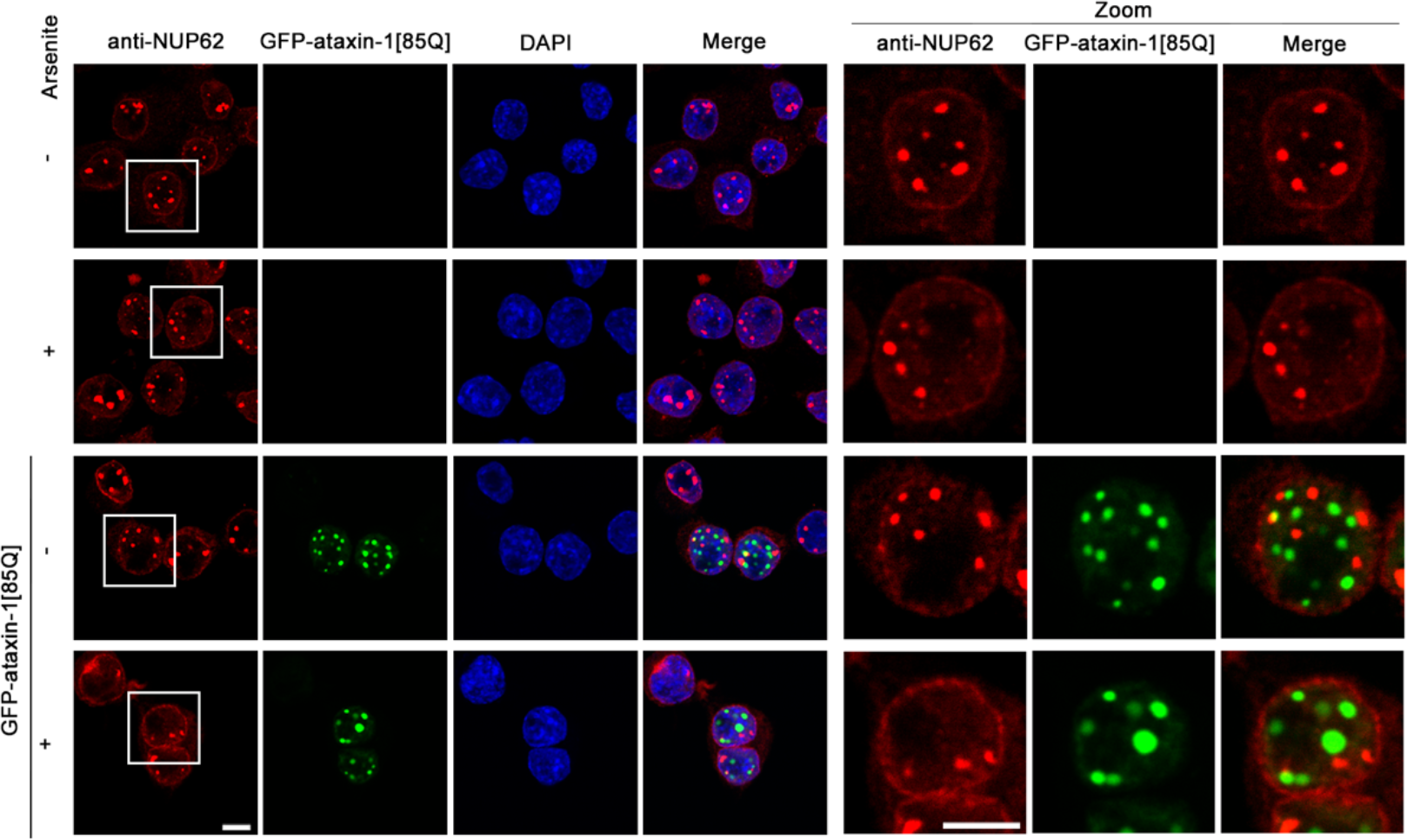
NUP62 localization is not altered by polyQ-ataxin-1 expression. Neuro-2a cells were transfected to express GFP-ataxin-1[85Q]. At 24 h post-transfection., cells were treated with arsenite as indicated, and then fixed, stained with anti-NUP62 and DAPI, before CLSM imaging. Representative images from 2 independent experiments are shown; merge panels overlay NUP62, GFP, and DAPI images. Zoom images (right panels) correspond to the boxed region. Scale bar = 10 μm.

**Figure S6.**
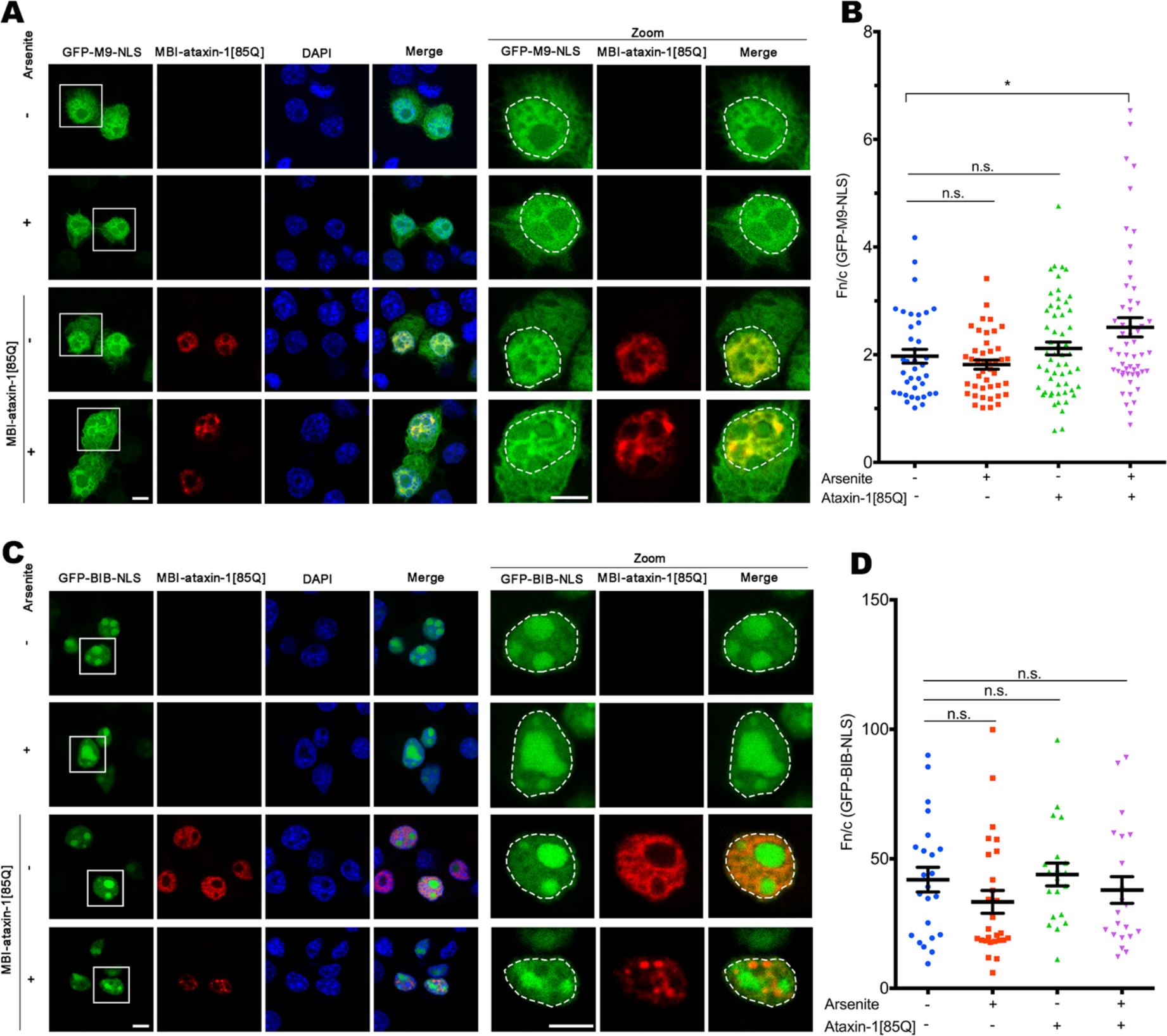
M9 NLS nuclear trafficking, but not BIB NLS nuclear trafficking, is disrupted in the presence of polyQ-ataxin-1. Neuro-2a cells were transfected to coexpress MBI-ataxin-1[85Q] with GFP-M9 NLS (A,B) or GFP-BIB (C,D). At 24 h post-transfection, cells were treated with arsenite as indicated, and then fixed, stained with anti-myc-antibody and DAPI, before CLSM imaging. (A & C) Representative images are shown from 3 independent experiments and merge panels overlay GFP, myc, and DAPI images; the position of the nucleus as determined by DAPI staining is indicated by the white dashed lines. Scale bar = 10 μm. (B & D) The nuclear to cytoplasmic fluorescence ratio (Fn/c) was calculated by a modified CellProfiler pipeline as per the legend to Figure 4B. Results represent the mean ± SEM (n > 25); * p<0.05, n.s. not significant.

**Supplementary Table S1: Alphabetical listing of the ataxin-1[85Q] interactome identified in Neuro2A cells by BioID and GFP-pulldown approaches**

(see separate Excel spreadsheet: Zhang_etal_SUPPTable1.xlsx)

**Supplementary Table S2:** Nuclear transporters (IPA: RAN Signaling) identified as members of the ataxin-1[85Q] interactome.

| **Uniprot ID_MOUSE** | **Encoded protein<sup>a</sup>** | **Nuclear Transport Role** |
| --- | --- | --- |
| IMA1 | Importin-α2 | Nuclear import receptor (α-family) binding classical NLS-containing proteins; heterodimerises with importin β1 to facilitate transit through the nuclear pore |
| IPO5 | Importin-5/<br>Importin-β3 | Nuclear import receptor (β-family) for proteins including some ribosomal proteins and histones |
| TNPO1 | Transportin-1/Importin-β2 | Nuclear import/export receptor (β-family) for proteins including hnRNPs, ribosomal proteins and histones |
| XPO1 | Exportin-1/CRM1 | Nuclear export receptor for diverse protein cargoes containing classical leucine-rich NES motifs |
| XPO2 | Exportin-2/CAS | Nuclear export receptor for importin-α |

<sup>a</sup> Includes alternative names

